# Whole genome sequence analysis reveals broad distribution of the RtxA type 1 secretion system and four novel type 1 secretion systems throughout the *Legionella* genus

**DOI:** 10.1101/768952

**Authors:** Connor L. Brown, Emily Garner, Guillaume Jospin, David A. Coil, David O. Schwake, Jonathan A. Eisen, Biswarup Mukhopadhyay, Amy J. Pruden

**Affiliations:** Via Department of Civil and Environmental Engineering, Virginia Tech, Blacksburg, VA 24061, USA; Department of Biochemistry, Virginia Tech, Blacksburg, VA 24061, USA; Department of Civil and Environmental Engineering, West Virginia University, Morgantown, WV 26506, USA; Genome Center, University of California, Davis, CA 95617, USA; Department of Natural Sciences, Middle Georgia State University, Macon, GA 31204, USA; Evolution and Ecology, Medical Microbiology and Immunology, University of California, Davis, CA 95167, USA

## Abstract

Type 1 secretion systems (T1SSs) are broadly distributed among bacteria and translocate effectors with diverse function across the bacterial cell membrane. *Legionella pneumophila*, the species most commonly associated with Legionellosis, encodes a T1SS at the *lssXYZABD* locus which is responsible for the secretion of the virulence factor RtxA. Many investigations have failed to detect *lssD*, the gene encoding the membrane fusion protein of the RtxA T1SS, in non-*pneumophila Legionella*, suggesting that this system is a conserved virulence factor in *L. pneumophila.* Here we discovered RtxA and its associated T1SS in a novel *Legionella taurinensis* strain, leading us to question whether this system may be more widespread than previously thought. Through a bioinformatic analysis of publicly available data, we classified and determined the distribution of four T1SSs including the RtxA T1SS and four novel T1SSs among diverse *Legionella* spp. The ABC transporter of the novel *Legionella* T1SS *Legonella* repeat protein secretion system (LRPSS) shares structural similarity to those of diverse T1SS families, including the alkaline protease T1SS in *Pseudomonas aeruginosa.* The *Legionella* bacteriocin (1–3) secretion systems (LB1SS-LB3SS) T1SSs are novel putative bacteriocin transporting T1SSs as their ABC transporters include C-39 peptidase domains in their N-terminal regions, with LB2SS and LB3SS likely constituting a nitrile hydratase leader peptide transport T1SSs. The LB1SS is more closely related to the colicin V T1SS in *Escherichia coli.* Of 45 *Legionella* spp. whole genomes examined, 19 (42%) were determined to possess *lssB* and *lssD* homologs. Of these 19, only 7 (37%) are known pathogens. There was no difference in the proportions of disease associated and non-disease associated species that possessed the RtxA T1SS (p = 0.4), contrary to the current consensus regarding the RtxA T1SS. These results draw into question the nature of RtxA and its T1SS as a genetic virulence determinant.

## INTRODUCTION

Type 1 secretion systems (T1SSs) are broadly distributed among bacteria and mediate the translocation of protein or peptide substrates with a broad range of function (1–4). The core T1SS complex is composed of a dimerized inner membrane ATP-binding-cassette (ABC) transporter protein, a trimerized membrane fusion protein that spans the periplasm, and a trimerized outer membrane protein. The genes encoding the ABC transporter and membrane fusion protein are typically localized together in the genome (5), while the gene encoding the outer membrane protein is usually encoded elsewhere, reflecting the multifunctional nature of this family of proteins (6,7).

Three major classes of T1SSs can be described based on the N-terminal region of the ABC transporter (4,8) (Figure 1). The first class includes bacteriocin transporters, such as the colicin V system in *Escherichia coli* (9), which encode ABC transporter proteins with N-terminal C-39 peptidase domains that cleave N-terminal regions of nascent substrates during translocation (4) (Figure 1A). In another class, such as the HlyA secretion system in *E. coli* (10), the ABC transporters contain an N-terminal C-39 peptidase-like domain (CLD), which lacks the catalytic histidine (9) (Figure 1B). A third class of T1SSs are composed of ABC transporters that lack either the C-39 peptidase or CLD. These systems typically secrete smaller substrates, including epimerases and proteases in *Azotobacter vinelandi* and *Pseudomonas aeruginosa*, respectively (11,12) (Figure 1C).

**Figure 1.**
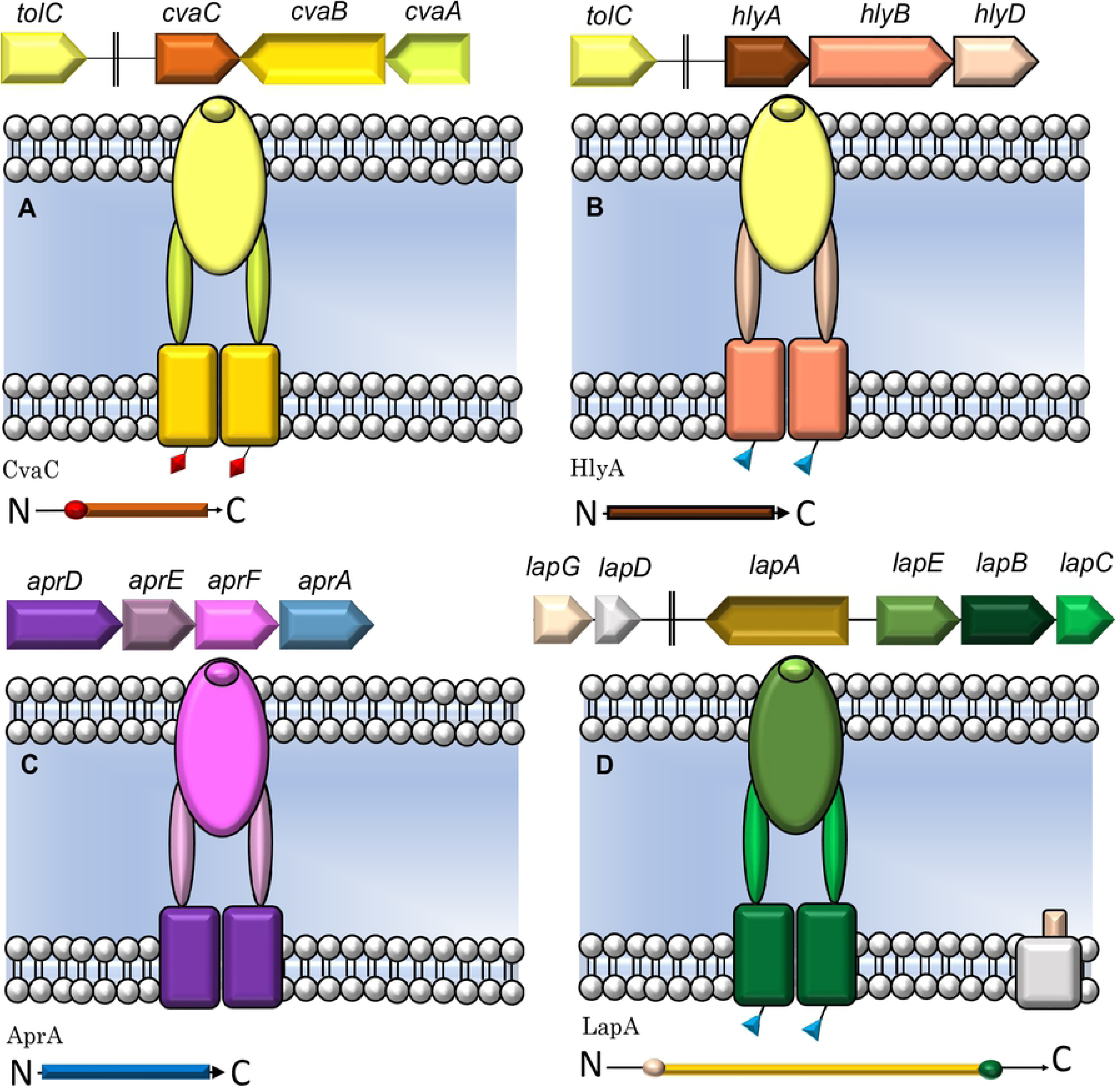
Genomic organization and structure of three type 1 secretion systems with corresponding substrates below each system [3, 8, 9]. (A) The CvaC T1SS in *E. coli.* An N-terminal region of CvaC is cleaved during secretion by the C-39 peptidase motifs on CvaB (red diamonds). (B) The HlyA T1SS in *E. coli.* HlyB has the CLD domain (blue triangles). (C) The AprA T1SS. AprD lacks either the CLD or C-39 peptidase motif. (D) The LapA T1SS in *P. fluorescens* Pf01. LapA is secreted in a two-step fashion where the retention module (tan circle on the N-terminus of LapA) is cleaved by LapG before release into the extracellular environment.

In *Legionella pneumophila*, the prototypical T1SS is encoded at the *lssXYZABD* locus with *lssB* and *lssD* encoding the ABC transporter and membrane fusion protein, respectively (13,14). This complex is responsible for secreting the virulence factor RtxA, which is associated with adherence, pore-formation, cytotoxicity, and entrance into host cells (15, 16). The RtxA T1SS belongs to a subset of the CLD-type T1SSs whose substrates possess an N-terminal retention module. The most well characterized system of this type is the adhesin LapA in *Pseudomonas fluorescens* strain Pf01 (3,17). In *P. fluorescens*, LapA is secreted in a two-step fashion mediated by membrane fusion protein LapC, the ABC transporter LapB, outer membrane protein LapE, and a transglutaminase-like cysteine proteinase (BTLCP) LapG (3,17–19) (Figure 1D). However, no study has detected *lssD* (a *lapE* homolog) in any non*-pneumophila* genome since the discovery of the *lssXYZABD* locus in 2001 (14,16,20). These observations have led to the assumption that RtxA is a key virulence determinant unique to *L. pneumophila*. In the present study, we report the broad distribution of the RtxA T1SS and three novel T1SSs throughout the *Legionella* genus and examine their occurrence among strains of disease-associated *Legionella*.

## RESULTS

### Isolation and phylogenetic identification of a novel *Legionella taurinensis* strain containing four type 1 secretion systems

During a survey of municipal and well waters in Genesee County, Michigan, endemic *L. pneumophila* were targeted for isolation using standard culture methods, and isolates were subjected to whole genome sequencing (Garner et al., *in review*). Sequence and phylogenetic analysis revealed the four isolates obtained from municipal water sourced from an aquifer to be novel *Legionella taurinensis* strains (21), a species that did not have a reference genome at that time (Table S1). These genome sequences are deposited at DDBJ/ENA/GenBank under the accession PRJNA450138. While analyzing the genomes of the isolates for virulence factors, the strain was found to possess the *lssXYZABD* locus believed to be absent from non-*pneumophila Legionella* spp. (14,16,20,22) in addition to three novel T1SSs with low homology to the RtxA T1SS components.

### Organization of the *lssBD* system in *Legionella* spp

In contrast to previous reports (14,20), *L. taurinensis* was found to possess the entire *lssXYZABD* locus associated with RtxA secretion with two reading frames (ORFs) (472 base pair and 4.24 kilobase) between *lssB* and *lssA*, both of which encode hypothetical proteins (Figure 2A, B, Figure S1, S2). *L. taurinensis* and *L. pneumophila* LssB possess the LapB-type N-terminal CLD (Figure S2, Table S2). *L. taurinensis* possesses the *lssXYZABD* locus including a gene encoding an LssD homolog which is 60% identical to *L. pneumophila* LssD (Figure S1).

**Figure 2.**
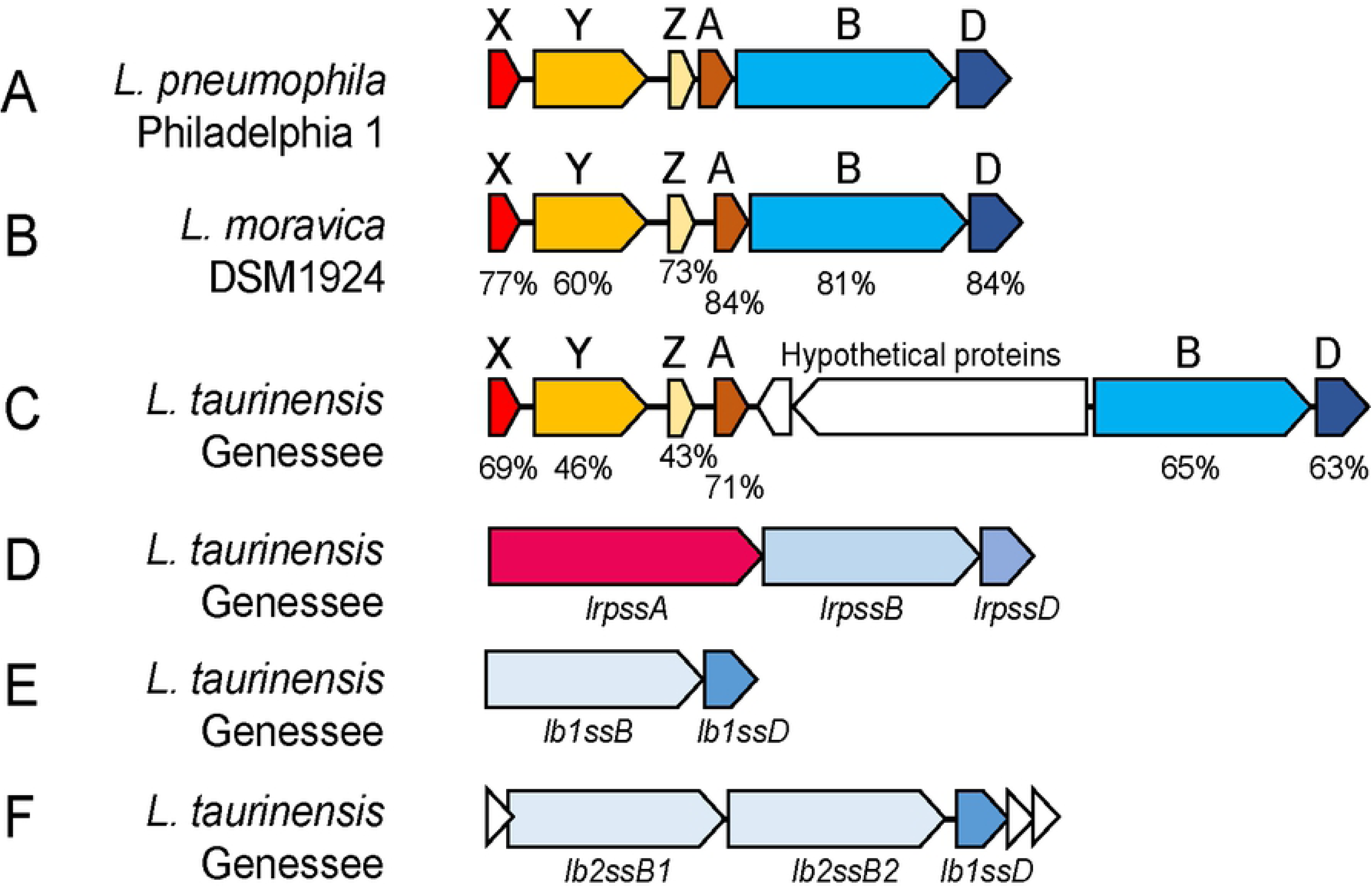
Genomic organization of four type 1 secretion systems in the *Legionella* genus. (A) Organization of the *lssXYZABD* locus in *L. pneumophila.* (B) The *lssXYZABD* locus in *L. taurinensis* encodes homologs of all members of the *lssXYZABD* locus, but the operon has two inserted ORFs. (C) The *lssXYZABD* locus in *L. moravica* DSM1924 (D) The *lrpss* T1SS found in *L. taurinensis* encodes a putative substrate at the LRPSS locus. (E and F) The *lb1ss* and *lb2ss* locus in *L. taurinensis.* White triangles are hypothetical proteins. (B-C) Percentages are percentage identity to *L. pneumophila* Philadelphia amino acid sequences. GenBank accession numbers are provided (Table S2).

Following this observation, we examined 45 *Legionella* species whole genome sequences for the RtxA T1SS by comparing amino acid sequences of *L. pneumophila* LssB and LssD against the predicted proteomes of *Legionella* spp. using blastp (Table S2). A species was considered to encode the RtxA T1SS if its genome encoded homologs of LssD and LssB with amino acid sequence ≥ 40% identity (Table S2) and if the ABC transporter was monophyletic with *L. pneumophila* LssB (Figure 3, Figure S3). These two proteins were chosen as they constitute two-thirds of the core components of a T1SS, the membrane fusion protein and ABC transporter. This analysis revealed that nearly half of species’ genomes examined (n = 19) encoded LssB and LssD homologs. Of these 19, several possessed the entire *lssXYZABD* locus, while others lacked some components of the locus or possessed additional ORFs within the cluster (data not shown). In all in which the RtxA T1SS was detected, *lssB* and *lssD* were encoded adjacent to one another in the genome.

**Figure 3.**
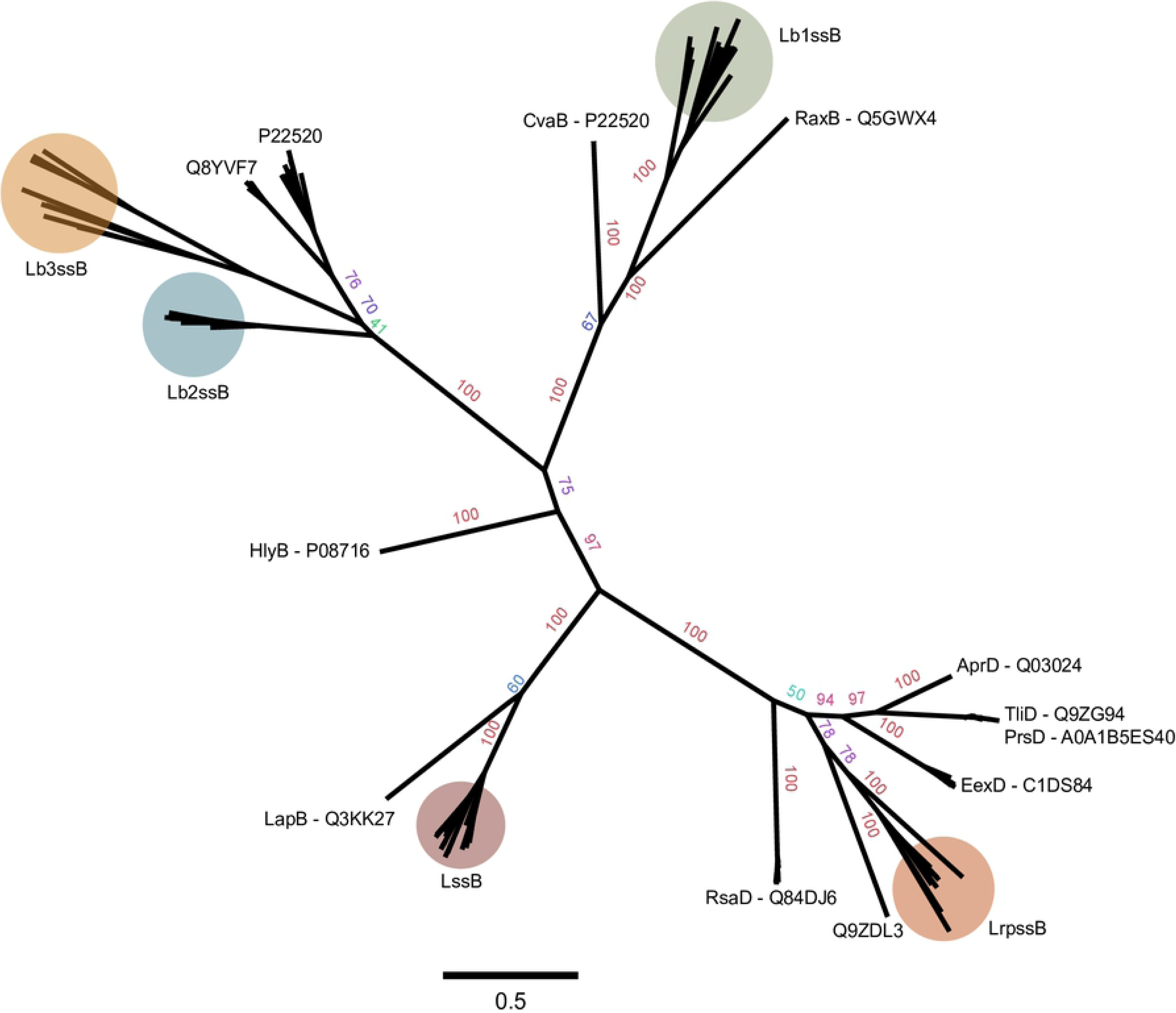
Unrooted maximum likelihood tree for trimmed sequences of 187 T1SS ABC transporters including the five *Legionella* systems. Bootstrap values displayed as percentages. An un-modified tree used to classify the *Legionella* secretion systems, and the *Legionella* sequences, are provided (Figure S5, Table S2).

**Table 1.** Results of tblastn searches against the *L. moravica* DSM19234 whole genome sequences. All hits had 99%-100% query coverage. Query proteins are all original protein sequences reported by *Jacobi et al* 2003 [15].

*Legionella moravica* strain DSM19234 was found to possess the entire *lssXYZABD* locus with corresponding proteins 60-86% identical to those encoded by *L. pneumophila* Philadelphia 1 (Figure 2A, C, Table 1). This is noteworthy as a previous bioinformatic investigation reported the absence of this locus in *L. moravica* DSM19234 in its entirety (22). This study reports using the tblastn algorithm (release 2.2.25) in BLAST to compare the protein sequences of the locus with nucleotide sequences of *Legionella* spp. Repeating this approach using the BLAST webserver against *L. moravica* DSM19234 whole genome sequences (taxid: 1122165), we detected all genes of the *lssXYZABD* locus with identity values ranging from 56.28%-84.13% (Table 1). Therefore, whole genome sequences of *L. moravica* DSM19234 were found to possess genes encoding the RtxA T1SS. Last, we additionally confirmed the presence of an RtxA-like substrate in a subset of genomes, including those of *L. taurinensis* Genessee01 and *L. moravica* species, by identifying T1SS substrates with a putative N-terminal di-alanine retention module specific to LapA/RtxA class substrates (3,17) through tblastn searches against their whole genome sequences (Figure S4). *Legionella longbeachae* is the second most commonly reported causative agent of Legionnaires’ Disease, especially in New Zealand and Japan where reported cases of *L. longbeachae* infection occur about as often as cases of *L. pneumophila* infection (23). Draft genome sequences of *L. longbeachae* strains F1157CHC and FDAARGOS-201 include *lssB* and *lssD* homologs with interrupting stop codons within the ORFs (Table S2) indicating that these genes are non-functional or contain sequencing or annotation errors. A tblastn search with LssB and LssD from *L. longbeachae* strain FDAARGOS-201 as queries did not identify respective homologs in the genome of *L. longbeachae* strain NSW150 (E-value cut-off of 1E-5). Therefore, the only finished genome of *L. longbeachae*, strain NSW150, lacks homologs of both *lssB* and *lssD.* Further, all three genomes of *L. longbeachae* were examined for homologs of the *L. pneumophila* Philadelphia 1 RtxA (lpg0645) and LapE (lpg00827) using tblastn and neither were detected (E-value cut-off 1E-5).

### *L. taurinensis* encodes three novel type 1 secretion systems

Analysis of the diversity of T1SSs across the *Legionella* genus has not been previously reported. Scanning the genome of *L. taurinensis* Genessee01 for additional *lssB* and *lssD* family genes revealed the presence of three novel T1SSs. One is composed of a 438 amino acid (aa) membrane fusion protein and 587 aa ABC transporter which were encoded adjacent to one another in the genome (Figure 2D). This ABC transporter lacks either the C-39 peptidase motif or CLD (Figure S5, S6) and a protein phylogeny of the ABC transporter indicated a relationship with T1SSs that secrete substrates of diverse function (Figure 3, Figure S3). Additionally, the genomic locus in *L. taurinensis* that encodes the ABC transporter and membrane fusion protein also encodes a putative substrate with hemolysin type calcium binding motifs commonly associated with T1SS substrates. Because of this, the name *Legionella repeat protein secretion system* (LRPSS) is proposed for this T1SS.

*L. taurinensis* additionally encodes two putative bacteriocin transport T1SSs. One is a colicin V-like T1SS (Figure 2F, Figure 3), for which the name *Legionella bacteriocin* 1 *secretion system* (LB1SS) is proposed. The second putative bacteriocin transporter locus encodes two ABC transporter proteins at the same genomic locus, (Figure 2E, Figure 3, Figure S5, Table S2), with one of these (PUT41641.1) including a C-39 peptidase motif at the N-terminus. This ABC transporter was found to be phylogenetically related to a nitrile hydratase leader peptide (NHLP)-type bacteriocin ABC transporter in *Nostoc sp. PCC 7120* (24) (Figure 3, Figure S3, Figure S5). For this system, the name *Legionella bacteriocin* 2 *secretion system* (LB2SS) is proposed. Additionally, searching the genomes of the *Legionella* spp. for homologs of the LB2SS system genes revealed a similar NHLP-type bacteriocin T1SS. This was suggested by protein phylogeny (Figure 3, Figure S3) to be a different T1SS or a highly diverged variant of the LB2SS. The name *Legionella bacteriocin 3 secretion system* (LB3SS) is proposed for this system.

### Distribution of T1SSs in the *Legionella* genus and their association with disease

We found that 19 of 45 (42%) *Legionella* spp. examined encode the RtxA T1SS (Figure 4). Of these 19, 7 (37%) are known pathogens. Relatively few (10 of 45, 22%) encode the LRPSS system, while 16 of the 45 species (35%) encode the LB1SS; 9 of 45 species (20%) encode the LB2SS, and 11 of 45 species (24%) encode the LB3SS. Collectively, only eight species lacked the T1SSs described in this paper. We performed significance testing because previous studies have qualitatively evaluated differences in the distribution of T1SSs in relation to perceived virulence or propensity for intracellular growth. Two proportion Z-tests with continuity correction were performed to compare the prevalence of the RtxA T1SS within strains who have never been isolated from patients and those which have (Figure 4). No significant differences were detected (p = 0.4), but the sample sizes are small (n = 19). Thus, there was no detected difference in the proportions of disease and non-disease associated species that possessed any of the T1SSs to the extent of our current knowledge of pathogenicity in these *Legionella* species. However, increasing the number of genomes analyzed could impact the results of the present analysis.

**Figure 4.**
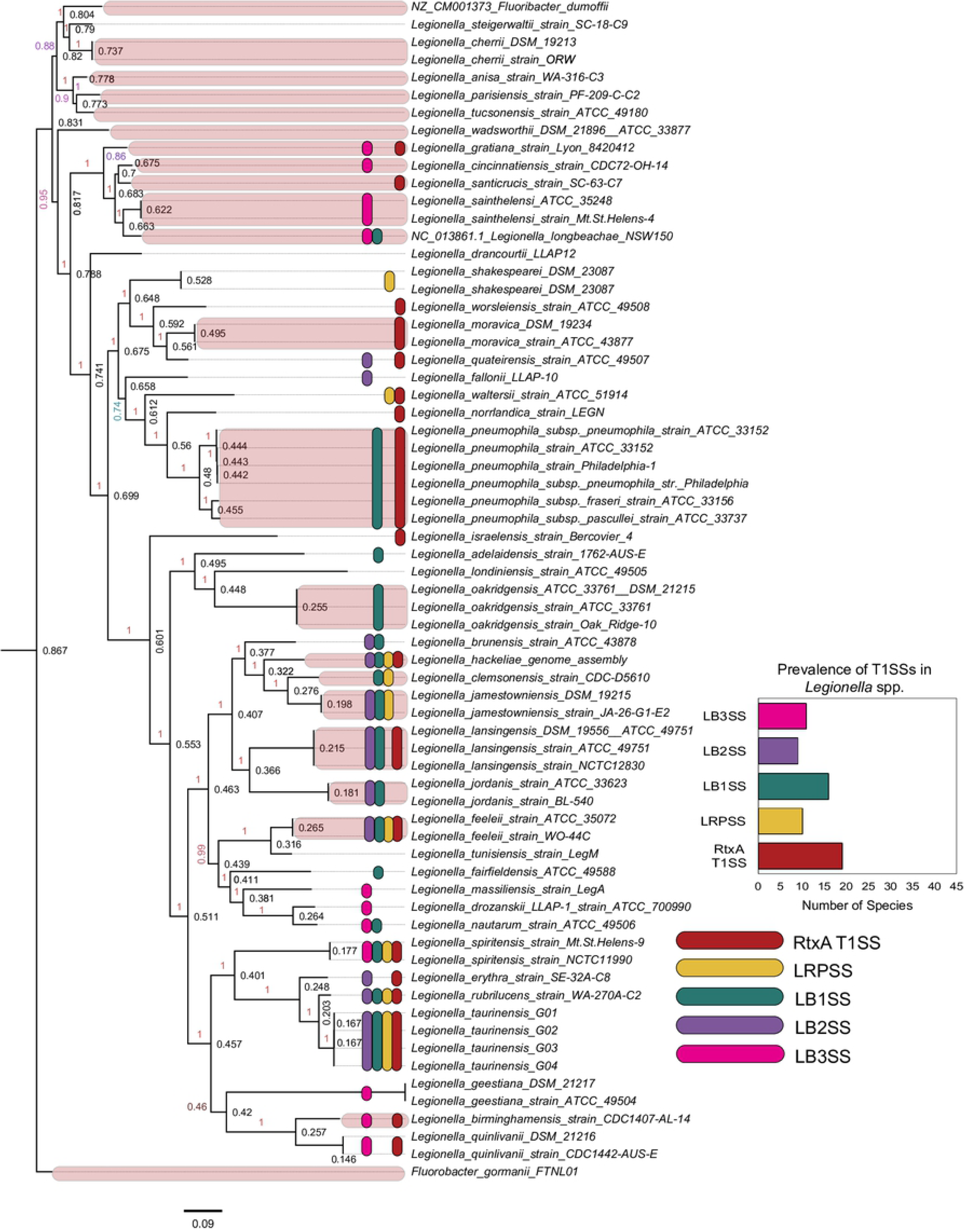
Nucleotide whole genome sequence FastTree tree predicted using a core set of marker genes predicted by PhyloSift of *Legionella* spp. overlaid with the distribution of T1SSs.

## DISCUSSION

Multiple investigations report the restriction of the *lssBD/*RtxA system to *L. pneumophila.* Two studies did not detect *lssD* in non-*pneumophila Legionella* using Southern blotting and DNA macroarrays, respectively (14,20), while one study detected *lssB* in all non-*pneumophila Legionella* tested (n = 10). Incidentally, the presence of *rtxA* (determined by Southern blotting) in *Legionella feeleii* has been reported, but the study did not test for the presence of the T1SS components (16). In retrospect, it may be unsurprising that several early studies did not detect the presence of *lssD* in non-*pneumophila Legionella* spp. given their reliance on DNA-DNA hybridization methods (14,15,20). This component may be more variable due to the nature of its interactions with a rapidly evolving substrate, and therefore methods relying on the nucleotide sequence of *L. pneumophila lssD* as probe could plausibly cause false-negatives of this nature. On the other hand, *Qin et al* 2017 bioinformatically examined non-*pneumophila Legionella* genomes (n = 21) for the *lssXYZABD* components (22) and reported the absence of the *lssXYZABD* locus from all strains tested, including *L. moravica* strain DSM19234. Further, this study reported that *Legionella* lacking the *lssXYZABD* locus displayed reduced intracellular multiplication relative to *L. pneumophila* strains which possess the T1SS (22). Thus, the emergent consensus model has been that the RtxA system is an important conserved genetic virulence determinant unique to *L. pneumophila.* In contrast, we found the RtxA T1SS and the three novel T1SSs to be prevalent throughout the genus, even among species not presently known to cause disease.

The 2014-2015 Center for Disease Control (CDC) Legionnaires’ Disease Surveillance Summary Report (25) documents the species isolated in culture confirmed Legionnaires’ Disease cases in the United States. Of 307 culture-confirmed cases in 2014-2015, 186 (60.6%) were caused by *L. pneumophila*, 3 (1%) by *L. longbeachae*, 4 (1.3%) by *L. micdadei*, 1 (0.3%) by *L. bozemanii* and 3 (1%) by *L. feeleii.* One hundred and eight (35%) were due to unreported species of *Legionella.* Of these, only *L. feeleii* and possibly some strains of *L. longbeachae* possess the RtxA T1SS. Additionally, *Gomez-Valero* et al 2019 measured replicative capacity of different *Legionella* spp. in THP-1 macrophages and found that *L. jordanis, L. taurinensis, L. jamestowniensis, L parisiensis, L brunensis*, and *L. bozemanii* displayed replicative capacity similar or superior to that of *L. pneumophila* (26). Of these species, *L. taurinensis, L. brunensis*, and *L. jamestowniensis*, (three of six (50%)) encode the RtxA T1SS. Therefore, while many *Legionella* spp. possess the T1SS responsible for RtxA translocation, our results indicate that the system alone does not predict propensity for disease association or intracellular replication in macrophages.

In conclusion, we report that the RtxA T1SS and four novel T1SSs discovered in *L. taurinensis* are broadly distributed among *Legionella* species and provide the first extensive survey and classification of T1SSs in the *Legionella* genus. Sequence and phylogenetic analysis indicate that *Legionella* spp. encode T1SSs that facilitate diverse functions, including bacteriocin transport and protein secretion. Importantly, no relationship was detected between the possession of the RtxA T1SS with disease association, drawing into question the nature of RtxA and its T1SS as a conserved genetic virulence determinant.

## Methods

### Determining Epidemiological Features of *Legionella* spp

We considered a *Legionella* species to be “disease-associated” based on whether not any strain of the species has ever been isolated from a patient. To determine this, we referenced the online resource at https://www.specialpathogenslab.com/legionella-species.php (accessed late 2018 - mid 2019) (27) and performed a literature review on PubMed Central to confirm accuracy.

### Sampling and culture Methods

Two one-liter samples of water were collected from one tap of a Genesee County School serviced by a groundwater well in March of 2016 into sterile polypropylene bottles (Nalgene, Rochester, NY) with 24 mg of sodium thiosulfate per liter added to quench chlorine for preservation prior to microbial analysis. Within 24 hours, samples were filter-concentrated onto a sterile 0.22 μm pore size mixed-cellulose ester membrane (Millipore, Billerica, MA) and resuspended in 5 mL sterile tap water prior to culturing according to International Standards of Organization (ISO) methods (28) for the recovery of *L. pneumophila* on highly selective buffered charcoal yeast extract media with supplemented glycine, polymyxin B sulfate, cycloheximide and vancomycin.

### Whole genome sequencing, assembly, and annotation

DNA was extracted using FastDNA SPIN Kit (MP Biomedicals, Solon, OH) according to manufacturer instructions. Purified DNA was quantified via a Qubit 2.0 Fluorometer (Thermo Fisher, Waltham, MA) and analyzed via gel electrophoresis to verify DNA integrity. Sequencing was conducted by MicrobesNG (Birmingham, United Kingdom) on a MiSeq platform (Illumina, San Diego, CA) with 2 x 250 bp paired-end reads. Libraries were constructed using a modified Nextera DNA library preparation kit (Illumina, San Diego, CA). Reads were trimmed using Trimmomatic (29) and *de novo* assemblies were generated using SPAdes (30). This Whole Genome Shotgun project has been deposited at DDBJ/ENA/GenBank under the accession PRJNA453403. The version described in this paper is version PRJNA453403.

### Sequence and phylogenetic analysis

The initial bioinformatic analysis that detected the T1SSs was performed using the integrated microbial genomes database and comparative analysis system (IMG) from the Joint Genome Institute (31).

To detect the RtxA T1SS components, *L. pneumophila* LssB and LssD amino acid sequences were compared with the protein sequences of *Legionella* spp. using BLASTp. For the three novel T1SSs, *L. taurinensis* amino acid sequences were used as query. A BLAST result was used to support a gene name when the amino acid sequence had overall identity of ≥40% (adjusted for incomplete query cover) with one of the query sequences. This criterion was used for all genes, except *lb1ssD*, for which many species displayed <40% amino acid homology relative to *L. taurinensis lb1ssD* (Table S2). Despite this, these genes were consistently found to be co-localized with *lb1ssB* homologs with ≥40% homology to *L. taurinensis* (for instance, WP_012979428.1/WP_012979429.1 in *L. longbeachae* strain NSW150). Last, protein phylogeny of the ABC transporters was inferred to validate the results suggested by the BLAST results. This analysis resulted in the renaming of several T1SSs which were near 40% homologous (Table S2). *Legionella* T1SS sequences with suggested names based on the results of the protein phylogeny and the sequence identity analysis are compiled (Table S2).

For the maximum likelihood (ML) tree, 187 amino acid sequences of T1SS ABC transporters were used. The sequences of previously described non-*Legionella* T1SS ABC transporters were chosen based on a review of the literature, especially (3,4,24). These sequences were then compared with the non-redundant protein database (32) using blastp and the ten best hits for each non-*Legionella* T1SS ABC transporter were collected. These sequences and the *Legionella* sequences were then aligned using MUSCLE v3.8.31 (33) with default settings. The aligned sequences were then trimmed using the heuristic method for trimAl, which resulted in a 482 amino acid aligned region (available in online materials) (34). Maximum likelihood trees for the trimmed sequence alignments were created using the RAxML webserver (35) with default settings including the GAMMA model of rate heterogeneity, the LG amino acid substitution matrix, and automatic bootstrapping (bootstopping cut-off = 0.03). The most likely tree was overlaid with bipartition values of 200 bootstrap replicates at the command line using RAxMLHPC v8.2.4. The tree constructed from the trimmed sequences is displayed in Figure 3.

### Whole genome sequence tree

Reference genomes from 45 *Legionella* species were downloaded from NCBI (Table S3). A set of core marker genes were identified using PhyloSift (36) and hmmer (37) to build a multiple sequence alignment. This alignment was used by FastTree (38) to generate a phylogenetic tree.

### Availability of data and material

All data including all *Legionella* sequences identified in the present study are provided in the paper and its supplementary materials, and may be accessed online through FigShare at doi:10.6084/m9.figshare.9162161.v1, doi:10.6084/m9.figshare.9162146.v1, doi:10.6084/m9.figshare.9162152.v1, doi:10.6084/m9.figshare.9162161.v1. The Whole Genome Shotgun project has been deposited at DDBJ/ENA/GenBank under the accession PRJNA450138. The version described in this paper is version PRJNA450138.

## Acknowledgements

The authors acknowledge and appreciate the time and support provided by Drs. William Rhoads and Marc Edwards.

## Supporting information captions

**Table S1.** Sequence analysis of conserved loci from the novel *Legionella taurinensis* strain reveal similarity between the novel strains and previously sequenced loci in *L. taurinensis*.

**Supplementary Figure 1.** Alignment of membrane fusion proteins from type 1 secretion system (T1SSs) of diverse functions. *L. pneumophila* LssD is 60% identical to *L. taurinensis* LssD.

**Supplementary Figure 2.** Alignment of CLD-type type 1 secretion system (T1SS) ABC transporters, including *Legionella pneumophila* LssB and the identified LssB homolog in *L. taurinensis*.

**Supplementary Figure 3.** Uncollapsed RAxML tree (Figure 3 of the main text).

**Supplementary Figure 4.** Partial alignment of *L. pneumophila* Paris, *L. taurinensis* Genessee01, and *L. moravica* RtxA. The black box encloses the sequences of the retention module.

**Supplementary Figure 5.** Alignment of C-39 peptidase-type T1SS ABC transporters including the ABC transporters of two novel T1SSs found throughout the *Legionella* genus. Both putative bacteriocin transporter ABC transporters contain C-39 peptidase motifs (red boxes) consistent with previously characterized system.

**Supplementary Figure 6.** Alignment of T1SS ABC transporters which do not possess N-terminal functional domains, including LrpssB discovered in *L. taurinensis*

**Supplementary Table 2.** NCBI Sequence IDs and BLAST results of *Legionella* T1SS components. All percent identities are adjusted for incomplete query covers, i.e. % query cover multiplied by % identity.

